# Design of High Affinity Binders to Convex Protein Target Sites

**DOI:** 10.1101/2024.05.01.592114

**Authors:** Wei Yang, Derrick R. Hicks, Agnidipta Ghosh, Tristin A. Schwartze, Brian Conventry, Inna Goreshnik, Aza Allen, Samer F. Halabiya, Chan Johng Kim, Cynthia S. Hinck, David S. Lee, Asim K. Bera, Zhe Li, Yujia Wang, Thomas Schlichthaerle, Longxing Cao, Buwei Huang, Sarah Garrett, Stacey R Gerben, Stephen Rettie, Piper Heine, Analisa Murray, Natasha Edman, Lauren Carter, Lance Stewart, Steve Almo, Andrew P. Hinck, David Baker

## Abstract

While there has been progress in the de novo design of small globular miniproteins (50-65 residues) to bind to primarily concave regions of a target protein surface, computational design of minibinders to convex binding sites remains an outstanding challenge due to low level of overall shape complementarity. Here, we describe a general approach to generate computationally designed proteins which bind to convex target sites that employ geometrically matching concave scaffolds. We used this approach to design proteins binding to TGFβRII, CTLA-4 and PD-L1 which following experimental optimization have low nanomolar to picomolar affinities and potent biological activity. Co-crystal structures of the TGFβRII and CTLA-4 binders in complex with the receptors are in close agreement with the design models. Our approach provides a general route to generating very high affinity binders to convex protein target sites.

## Main text

Naturally occurring high affinity protein-protein interfaces generally exhibit considerable shape complementarity, which enables concerted interatomic interactions and solvation free energy reduction needed to overcome the entropic cost of macromolecular association.^1^ To design proteins which bind to a target of interest with high affinity, it is similarly important that the shape of the designed binder and the target be complementary over the targeted region. There has been considerable recent progress in the design of small (50-65 residue) mini proteins to bind to targets of interest^2,3^, but since miniproteins in this size range are roughly spherical in shape and hence have convex binding surfaces, they are not well suited to bind to convex protein target sites due to the requirement for overall shape matching (Fig 1a,b). Methods for designing proteins which bind to convex target sites could considerably expand the power and scope of *de novo* binder design.

**Figure 1.**
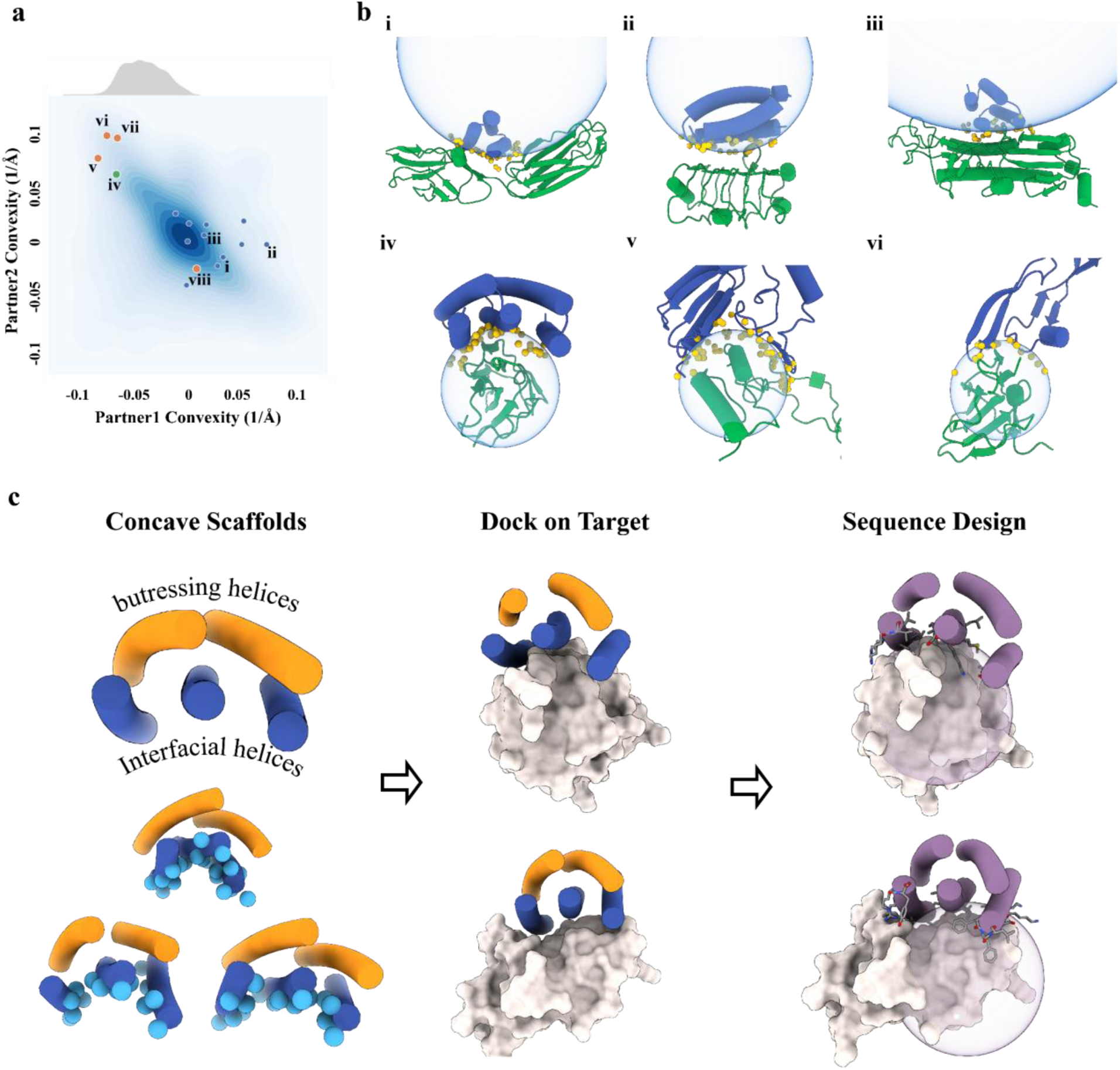
Design of 5HCS scaffolds to target convex interfaces. **a**, Distribution of protein-protein interface curvatures from the PDB and designed protein binders. Blue dots: previously designed protein binders (for these, the designed binders are partner 1 and the targets, partner 2). Previously designed protein binders^3^ have been limited to binding to flat or concave interfaces (receptor convexity <=0). Orange dots: examples of native protein complexes, v: PDB ID, 5XXB; vi: TGFβIII/TGFβRII complex, PDB ID, 1KTZ, vii: CD86/CTLA-4 complex, PDB ID, 1I85, viii, PD-1/PD-L1,PDB ID, 3bik. The TGFβRII and CTLA-4 functional interfaces showed high convexity, which we used as case studies to design concave binders. Green dot: The 5HCS scaffolds described in this paper can target convex binding sites. The distribution of convexity of the 5HCS scaffolds (upper part of panel a) shows that the 5HCS scaffolds are diverse enough to cover most of the naturally existing convex interfaces. **b.** Design models of complexes highlighted in panel a. i,ii,ii are PDGFR, 1GF1R, H3 in complex with corresponding *de novo* minibinders; iv, 5HCS binder in complex with TGFβRII; v, PDB ID: 5XXB; vi, TGFβIII/TGFβRII complex, PDB ID:1KTZ. Binders and receptors are shown as blue and green cartoons, respectively. Interfacial heavy atoms from binders are shown as yellow solid spheres. Fitted spherical surfaces are shown as blue transparent spheres. **c,** Design workflow. Column 1: 5HCS concave scaffolds with a wide range of curvatures were designed with three helices (blue) forming the concave surfaces (Cbeta labeled as spheres) and two helices (orange) buttressing at the back side. Column 2: Docking of 5HCS scaffolds to target binding sites. Column 3: Following docking, the interface sequencing is optimized for high affinity binding.

We reasoned that to enable systematic design of binders to convex protein target sites it would be necessary to generate scaffold sets with overall concave shapes. Three additional properties would further facilitate binder design and characterization. First, varying curvature: protein surfaces vary considerably in shape, and to enable close complementary matching of a wide range of targets, a set of proteins with varying curvature and surface topography would be ideal. Second, high stability: the higher the stability of the base scaffold, the more room for customizing the binding interface for high affinity binding, and the more robust the resulting binders. Third, minimal size: to make design cost effective for gene synthesis and generation of oligonucleotide libraries for initial screening, and for applications such as tumor penetration in oncology, the overall length of the binder scaffolds should be minimal (80-120 aa). With these selection criteria, we set out to construct a set of scaffolds.

## Results

### Computational design of 5HCS scaffolds

Previous work with designed helical repeat proteins (DHRs) has demonstrated that a wide range of curvatures can be obtained while maintaining high stability, but these proteins are generally well over 150 residues in length with 8 or more helices^4,5^. We reasoned that scaffolds resembling DHRs but with fewer helices could satisfy the four target properties (concave interaction surface, tunable curvature, high stability, and minimal size). We focused on 5 helix bundle scaffolds with three helices forming the concave interface and two helices providing structural support (Fig. 1c). Scaffolds were generated using a library of ideal helical and loop fragments combined first to create helix-turn-helix-turn modules. The length of each helix was constrained to 18 to 22 amino acids (5 to 6 helical turns) balancing stability and overall length constraints. These modules were repeated 3 times to generate 3 unit repeat proteins, and either the N-or C-terminal helix was truncated to generate five helix proteins with fewer than 120 amino acids. We evaluated the curvatures of the surfaces formed by the three interfacial helices and filtered out the backbones with convex surfaces. To more extensively diversify the backbones and break the repeat symmetry, we remodeled the backbones by randomly replacing short scaffold structural elements with alternative local structures (fragments) from the PDB followed by Cartesian-space minimization and full-atom optimization^6^. Following Rosetta sequence design^7^, designs with sequences predicted to fold to the designed structure with AlphaFold2^8^ and to have high accuracy using DeepAccNet^9^ were selected (Fig. S1). The selected 7476 scaffolds, which we refer to as 5HCS (5 helix concave scaffolds) throughout the remainder of the text, have a wide range of curvatures (Fig. S1).

To guide selection of representative convex targets for 5HCS scaffolds, we systematically analyzed the convexities of protein-protein interfaces from the PDB. Pairs of interacting proteins were extracted from PDB entries with multiple chains and were grouped into 2411 clusters. We calculated the convexities of representatives of each cluster by fitting the interfacial heavy atoms to spherical surfaces using the random sample consensus (RANSAC) algorithm^10^. Because of the overall shape matching constraint, the convexity of the two binding partners for each complex are negatively correlated: when one partner is convex, the other is almost always concave (Fig. 1a). Previously designed mini binders^3^ are flat or convex, and bind to flat or concave targets (Fig. 1a,b). The convexities of the 5HCS scaffolds covers the range of convexities we analyzed from PDB (Fig. 1a).

### Design and structural validation of TGFβRII binders

We selected as a representative convex target site that of the transforming growth factor-β3 (TGF-β3) on the TGF-β receptor type-2 (TGFβRII) (PDB ID: 1KTZ). Binders to modulate TGF-β pathways have considerable interest as therapeutics in oncology, tissue fibrosis, and other areas^11^. We used the RIF based docking protocol of Cao et al^3^ to dock both the 5HCS scaffolds described above and the globular miniprotein scaffold library used in the previous studies^2,12^ to the TGF-β binding site^13^ (Fig. S6a). Following design and filtering^14^ for binders with the concave surface of the 5HCS interacting with the target, and Alphafold2^8^ based confirmation of structure and binding mode, we encoded the designs using oligonucleotide arrays and cloned into a yeast surface-expression vector to enable high throughput assessment of binding affinity. After two rounds of fluorescent activated cell sorting for binding to biotinylated TGFβRII, sequencing revealed 2 5HCS hits but no mini-protein hits despite the nearly 100-fold greater representation of the latter in the library (see Methods). The sequences of the two hits are identical to the two designed sequences. We further optimized the most enriched design 5HCS_TGFBR2_0, by resampling the sequences of interfacial residues in the bound state using ProteinMPNN^15^ and filtering the complex models using Alphafold2^8^. We combined the mutations predicted to improve binding affinities and encoded a combinatorial library with these mutations included using degenerate codons (Fig. S2a). Finally, we sorted the library for the optimized binders using yeast display selection (Fig. S3).

Four of the optimized binders obtained after several rounds of yeast display selection were produced in *E. coli*. The highest affinity binder, 5HCS_TGFBR2_1, was found using biolayer interferometry to have an affinity less than 1nM for TGFβRII (Fig. 2c, Fig. S5a). The sequence identity between 5HCS_TGFBR2_0 and 5HCS_TGFBR2_1 is 88.12% (Fig. S4a). The circular dichroism spectra indicates a helical structure with peaks at 208 nm and 222 nm consistent with the design model (Fig. 2a,b), and was only slightly changed by heating to 95 °C, indicating high stability (Fig. 2b).

**Figure 2.**
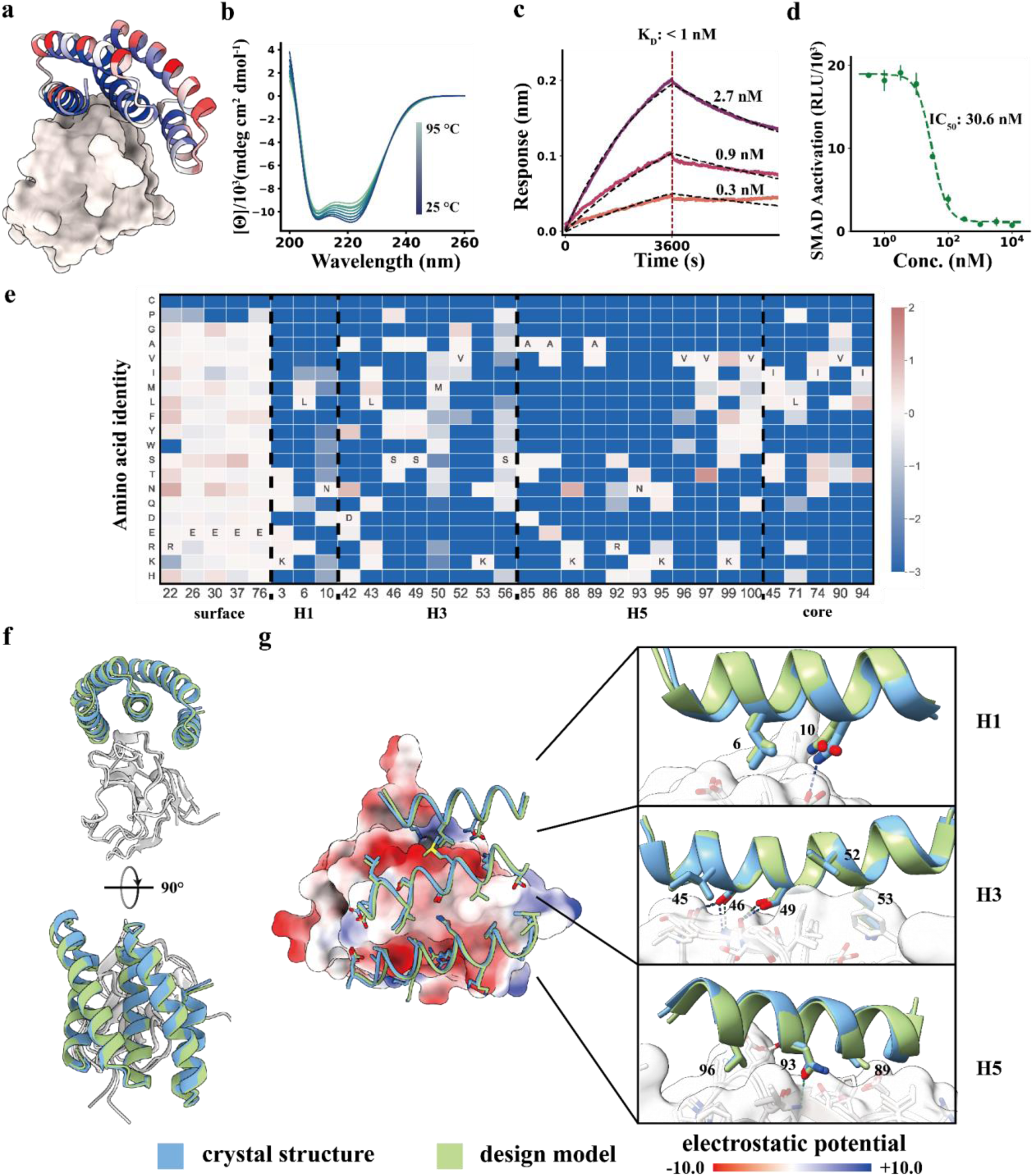
Concave 5HCS binder to TGFβRII. **a**. Design model of 5HCS_TGFBR2_1 (cartoon) binding to TGFβRII (PDB ID: 1KTZ). 5HCS_TGFBR2_1 is colored by Shannon entropy from the site saturation mutagenesis results at each position in blue (low entropy, conserved) to red (high entropy, not conserved). **b.** Circular dichroism spectra from 25 °C to 95 °C for 5HCS_TGFBR2_1. **c.** Biolayer interferometry characterization of 5HCS_TGFBR2_1. Biotinylated TGFβRII were loaded to Streptavidin (SA) tips and incubated with 2.7 nM, 0.9 nM and 0.3 nM of 5HCS_TGFBR2_1 to measure the binding affinity. The binding responses are shown in solid lines and fitted curves shown in dotted lines. **d.** Dose-dependent inhibition of TGF-β3 (10 pM) signaling in HEK293 cells. The mean values were calculated from triplicates for the cell signaling inhibition assays measured in parallel, and error bars represent standard deviations. IC_50_ values were fitted using four parameter logistic regression by python scripts. **e.** Heat map of the log enrichments for the 5HCS_TGFBR2_1 SSM library selected with 1.6 nM TGFβRII at representative positions. Enriched mutations are shown in red and depleted in blue. The annotated amino acid in each column indicates the residue from the parent sequence. **f,g.** Crystal structure of 5HCS_TGFBR2_1 in complex with TGFβRII. Left are top and side views of the crystal (blue and gray) superimposed on the design models (green and white). In the middle, TGFβRII is shown in surface view and colored by electrostatic potential (using ChimeraX; red negative, blue positive). On the right, detailed interactions between 5HCS_TGFBR2_1 (blue, green) and TGFβRII (gray, white) are shown.

We determined co-crystal structures of 5HCS_TGFBR2_1 with TGFβRII. The high resolution (1.24 Å) X-ray crystal structure is very close to the computational design model (Fig. 2f,g; root mean square deviation (rmsd) over C_α_ atoms of 0.55 Å over the full complex), showing 5HCS_TGFBR2_1 binds to the TGF-β3 binding site on TGFβRII utilizing the concave surface as designed.

To further investigate the sequence dependence of folding and binding, we generated site saturation mutagenesis (SSM) libraries in which each residue was substituted with all other nineteen amino acids one at a time, and sorted the library using fluorescence-activated cell sorting (FACS) with fluorescent TGFβRII. Deep sequencing results were closely consistent with the design model and crystal structure. Both the core residues and interfacial residues were highly conserved, while surface residues not at the interface were quite variable (Fig. 2 a,e). Helices H1, H3 and H5 which form the concave binding surface interact with TGFβRII, and the most highly conserved non-core residues are in these helices. In H1, N10 hydrogen bonds with TGFβRII D142 (Fig 2g, top panel); in H3, S46 and S49 hydrogen bond to the backbone atoms of strand S72 – S75 (Fig 2g, middle panel); and in H5, N93 hydrogen bonds to the backbone atoms of I76 (Fig 2g, lower panel). A hydrophobic patch composed of F48, L50 and I76 on TGFβRII critical for TGF-β3 binding packs tightly on a hydrophobic groove formed by L6 from H1, M50, V52, K53 from H3 and V96, K99, V100 from H5 (Fig. S7). All the key interactions described above are recapitulated in the crystal structure with high side-chain orientation consistency (Fig. 2f,g). Design of such extended grooves and pockets is nearly impossible using small globular miniproteins; the high affinity binding and crystal structure of 5HCS_TGFBR2_1 demonstrates that 5HCS scaffolds can indeed be used to target convex binding sites.

We assessed the biological activities of 5HCS_TGFBR2_1 in cell culture signaling assays. HEK293 cells with luciferase reporter for the TGFβ SMAD2/3 signaling pathway were stimulated using 10 pM TGF-β3 and varying concentrations of 5HCS_TGFBR2_1. Dose-dependent inhibition of the TGFβ SMAD2/3 signaling was observed with an IC_50_ of 30.6 nM (Fig. 2d).

### Design and structural validation of CTLA-4 binders

An important class of convex targets are the portions of the extracellular domains of transmembrane receptors which interact with their biological partners. These frequently consist of immunoglobulin fold domains, which our large scale shape analyses indicate are generally quite convex (Fig. S8). Immunoglobulin domain recognition plays important roles in immune receptor functions^16^; in particular the cancer immunotherapy targets Cytotoxic--T--lymphocyte--antigen--4 (CTLA-4) have extracellular Ig fold domains that are the targets of therapeutic antibodies^17^. Because of the therapeutic importance of the target, and receptor extracellular Ig domains more generally, we next sought to evaluate the generality of our approach by designing 5HCS based binders to CTLA-4.

CTLA-4 plays an important role in peripheral tolerance and the prevention of autoimmune disease by inhibition of T cell activation. Antibody CTLA-4 targeting checkpoint inhibitors^18^ have been used for melanoma and non-small cell lung cancer (NSCLC) therapy. We targeted the region surrounding the beta-turn (132-140) of CTLA-4 which is buried in the interface between CTLA-4 and CD86 (PDB ID: 1I85) (Fig. S9a) using the methods described above. FACS of yeast libraries displaying the concave designs identified six CTLA-4 binders. The sequences of the six hits match their designs with 100% sequence identity. Deep sequencing of a site saturation mutagenesis library of the most enriched binder, 5HCS_CTLA4_0, showed that the designed core and interfacial residues of the binder were highly conserved, suggesting the design folds and binds target as in the computational model (Fig. 3a,e, Fig. S9a). As the Alphafold2^8^ predicted models were not consistent with the designed complex model, we combined the most enriched substitutions from the SSM heatmap, instead of using the ProteinMPNN^15^ resampling followed by Alphafold2^8^ filtering method mentioned above.

**Figure 3.**
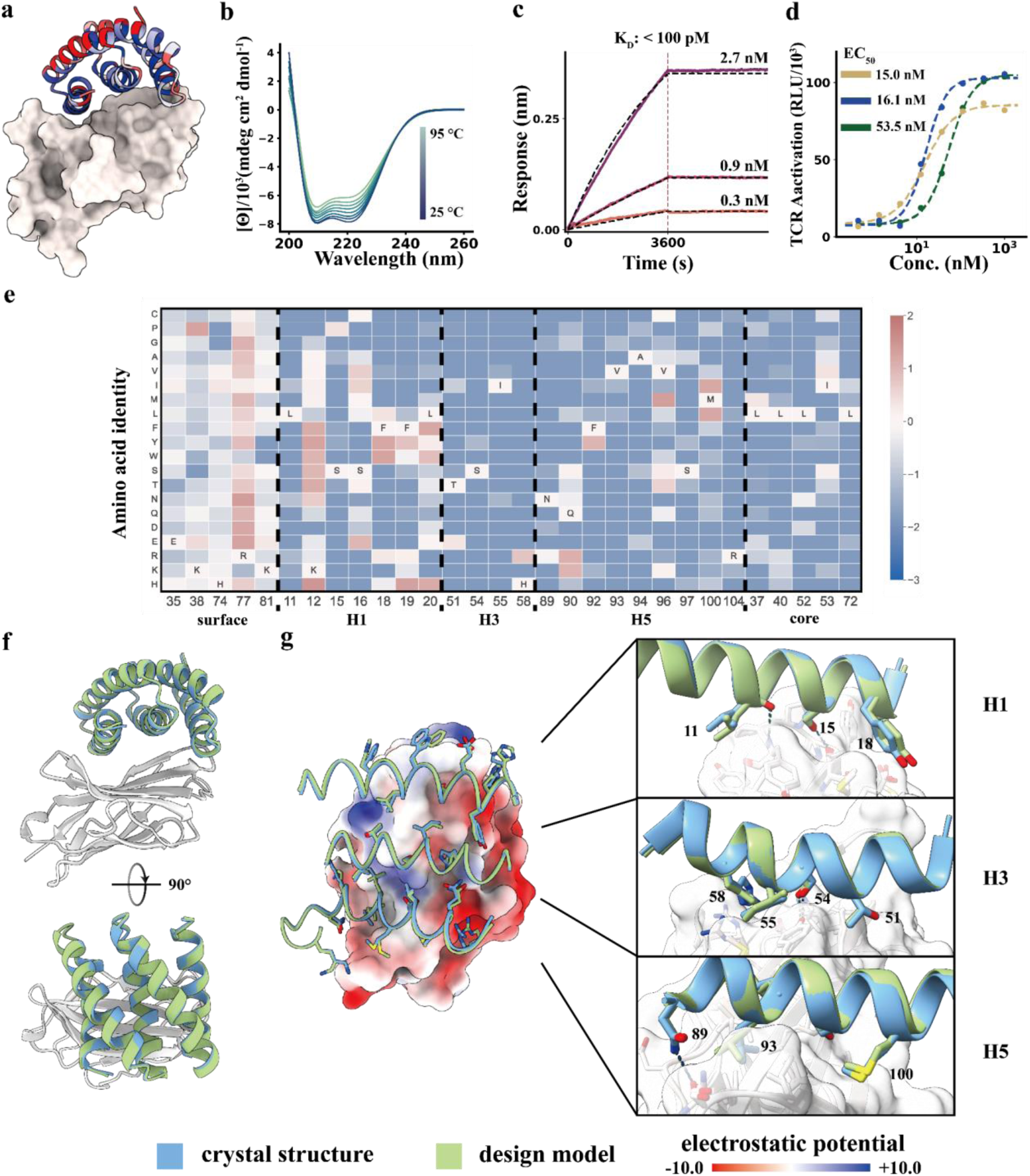
Designed 5HCS CTLA-4 binder. **a**. Model of 5HCS_CTLA4_1 (cartoon) binding to CTLA-4 (PDB ID:1l85) colored by Shannon entropy from site saturation mutagenesis results. **b.** Circular dichroism spectra from 25 °C to 95 °C for 5HCS_CTLA4_1. **c.** Biolayer interferometry characterization of 5HCS_CTLA4_1. Biotinylated CTLA-4 was loaded to Streptavidin (SA) tips and these were incubated with 2.7 nM, 0.9 nM and 0.3 nM of 5HCS_CTLA4_1 to measure the binding affinity. **d.** Increase of TCR activation induced signal (via NFAT pathway) from engineered CTLA-4 effector cells lines by 5HCS_CTLA4_1 (green), lpilimumab (gold) and 5HCS_CTLA4_1_c6 (blue) is shown. EC_50_ values were fitted using four parameter logistic regression by python scripts. Color schemes and experimental details are as in Fig 2. **f.** Designed interactions between 5HCS_CTLA4_1 (green) and CTLA-4 (white). **e.** Log enrichments for the 5HCS_CTLA4_1 SSM library selected with 10 nM CTLA-4 at representative positions. The annotated amino acid in each column indicates the residue from the parent sequence. **f,g.** Crystal structure of 5HCS_CTLA4_1 in complex with CTLA-4. Color schemes are the same as Fig. 2.

We synthesized the combinatorial library with these mutations included using degenerate codons (Fig. S2b). After additional rounds of yeast display selection of the combinatorial library, we expressed four of the best binders in *E. coli*. The highest affinity optimized binder, 5HCS_CTLA4_1 has a sequence similarity of 82.86% compared to 5HCS_CTLA4_0 (Fig. S4b). 5HCS_CTLA4_1 had an off rate too slow and a binding affinity for CTLA-4 too tight (<100 pM) to be measurable by biolayer interferometry (Fig. 3c, Fig. S5b).

We determined co-crystal structures of 5HCS_CTLA4_1 with CTLA-4 and unbound crystal structures of 5HCS_CTLA4_2 (Fig. S10, Table S2). The unbound crystal structure of 5HCS_CTLA4_2 aligns with the structure of binder in 5HCS_CTLA4_1 bound structure well with a rmsd. of 0.416 Å. The design model of 5HCS_CTLA4_1 in complex with CTLA-4 also closely agrees with the crystal structure, with a very low rmsd of 0.34 Å (Fig 3f,g). 5HCS_CTLA4_1 binds to the CD86 binding site on CTLA-4 using a concave binding surface formed by H1, H3 and H5 covering both the CTLA-4 beta-turn (L98 to Y104) and hydrophobic pocket which interacts with CD86 (Fig. S9b). H1 interacts with the hydrophobic beta-turn (L128 to Y136) through hydrophobic interactions between Y18 and M135 and aromatic interactions between H19 and Y136 (Fig. 3g, top panel). Substitution of this residue with H or Y improves binding affinity (Fig 3E). S54 and I55 on H3 interact with Y139 on CTLA-4 (Fig 3g, middle panel), and N89 on H5 hydrogen bonds with Q90 on CTLA-4. (Fig. 3g, lower panel). All of these interactions are closely recapitulated in the crystal structure (compare blue and green in Fig 3g). The circular dichroism spectra indicates a helical structure with peaks at 208 nm and 222 nm consistent with the design model and was unchanged by heating to 95 °C, indicating high thermal stability (Fig. 3b).

We tested the biological activity of 5HCS_CTLA4_1 in cell culture using an immune checkpoint functional assay in which stably expressing CTLA-4 Jurkat cells with a luciferase reporter for TCR/CD28 activation were incubated with activating Raji cells expressing the CTLA-4 ligands CD80 and CD86. Inhibition of the inhibitory CTLA-4 – CD86 interaction results in TCR pathway activation, and hence can be directly read out using this assay. We co-cultured the cells with a range of concentrations of the CTLA–4 binder, and observed dose-dependent activation of CTLA-4 effector cells with an EC_50_ of 53.3 nM (Fig. 3d). Surprisingly, this is higher than the EC_50_ (15.0nM) of the anti-CTLA-4 antibody lpilimumab (MDX-010, Yervoy), despite the at least two order weaker binding affinity for CTLA-4 (18.2 nM)^19^. Steric or avidity effects may contribute to the potency of the antibody, which can interact with two receptors through the two Fabs. To explore the effect of avidity, we flexibly fused 5HCS_CTLA4_1 to previously designed domains which oligomerize into different symmetric architectures^20^. We found that a highly expressed and monodisperse hexameric version (Fig. S11), 5HCS_CTLA4_1_c6 had an EC_50_ of 16.1 nM, comparable to the antibody (Fig. 3d).

### Design and structural validation of PD-L1 binders

Programmed death-ligand 1 (PD-L1), is upregulated on many tumors, and interacts with PD-1 on T-cells to downregulate T-cell activation. Therapeutic antibodies against PDL1 have shown considerable promise for checkpoint inhibition in cancer immunotherapy^18^. Considering the therapeutic importance of the target, and to test the generalizability of our approach towards flatter protein surfaces, we designed binders using the methods described above to target the binding site of PD-1 on PD-L1 (PDB ID: 3BIK) and block the interaction between the two proteins (Fig. 1a, Fig. S12a). Two PD-L1 binders were obtained from a set of 96 designs. We optimized the stronger binder, 5HCS_PDL1_0, by resampling the residues at the designed interface using ProteinMPNN^15^ followed by Alphafold2^8^ filtering. We used yeast display to sort a library with degenerate codons encoding mutations (Fig. S2c) predicted to improve binding, expressed ten of the most enriched binders in *E. coli*, and measured their binding affinities by biolayer interferometry. The highest affinity binder, 5HCS_PDL1_1, which has 93.2% similarity with the sequence of 5HCS_PDL1_0 (Fig. S4c), is expressed at high levels, very stable (Fig 4b) and has an affinity of 646 pM (Fig. 4c, Fig.S5c).

**Figure 4.**
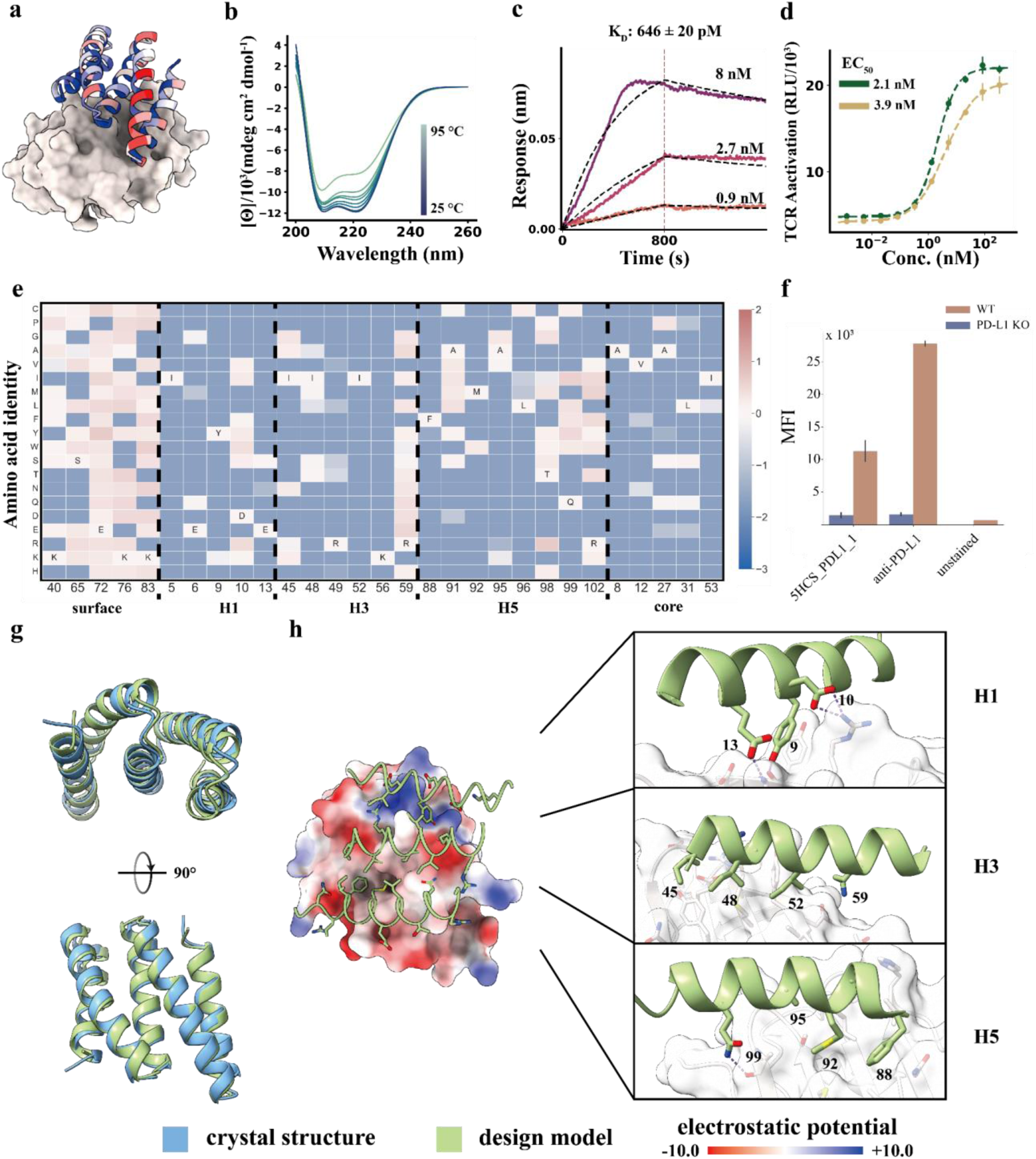
Designed 5HCS binder to PD-L1. **a**. Model of 5HCS_PDL1_1 (cartoon) binding to PD-L1 (PDB ID: 3BIK), with 5HCS_PDL1_1 colored by Shannon entropy from site saturation mutagenesis results. **b.** Circular dichroism spectra from 25 °C to 95 °C for 5HCS_PDL1_1. **c.** Biolayer interferometry characterization of 5HCS_PDL1_1. Biotinylated PD-L1 was loaded to Streptavidin (SA) tips and these were incubated with 8 nM, 2.7 nM and 0.9 nM of 5HCS_PDL1_1 to measure the binding affinity. **d.** The increase of TCR activation induced signal (via NFAT pathway) from engineered PD-1 effector cells lines by 5HCS_PDL1_1 (green), control antibody (gold) is shown. The mean values were calculated from triplicates for the cell signaling inhibition assays measured in parallel, and error bars represent standard deviations. Color schemes and experimental details are as in Fig3. **e.** Heat map representing the log enrichments for the 5HCS_PDL1_1 SSM library selected with 6 nM PD-L1 at representative positions. The annotated amino acid in each column indicates the residue from the parent sequence. **f.** WT A431 and PD-L1 KO A431 cell lines were stained with fluorophore labled 5HCS_PDL1_1 and anti-PD-L1 antibody respectively and then analyzed through FACS. **g,h.** Unbound crystal structure of 5HCS_PDL1_1 and designed interactions between 5HCS_PDL1_1 (green) and PD-L1 (white). Color schemes are the same as Fig. 2.

To examine the sequence determinants of folding and binding of 5HCS_PDL1_1 and to provide a structural footprint of the binding site, we generated a SSM library and sorted the library using FACS with fluorescent PD-L1. The conservation of both the core residues and the interfacial residues (Fig. 4a,e, Fig. S12c) suggests the binders fold and bind to the models as designed. As with 5HCS_TGFBR2_1 and 5HCS_CTLA4_1, the interfacial helices H1, H3 and H5 of 5HCS_PDL1_1 have an overall concave shape (Fig. 4a). The key interactions between H1 and PD-L1 include aromatic packing of Y9 and Y123 on PD-L1 and electrostatic interactions between D10 and E13 with K124 and R125 on PD-L1 (Fig. 4h). H3 binds to the hydrophobic pocket formed by Y56, M115, A121 and Y123 (Fig. 4h). Residues Y9, E13, K56 and Q99 spanning the three helices satisfy the hydrogen bonding requirements of both the side chains and backbone of the PD-L1 edge beta strand (A121-R125) buried at the interface.

We solved the high resolution crystal structure of 5HCS_PDL1_1. The refined structure has excellent geometry (Table S2) and reveals the expected helical assembly with five antiparallel helices (Fig. 4g,S10b). The crystal structure of 5HCS_PDL1_1 superimposes on the computational design model with a rmsd of 0.75 Å over 105 aligned Cα atoms (Fig. 4g, Fig. S10b; the substitutions which increase affinity relative to 5HCS_PDL1_0 do not alter the backbone structure). Not surprisingly, given the near identity between the computational designs (Fig. S4c) and the crystal structures, the shape and electrostatic potential of the designed target binding interfaces are nearly identical between the crystal structure and the computational design model (Fig. S10c).

### Comparison to DARPINs

While the computational design of extended concave binding proteins has not been possible to date, DARPIN binders based on the native ankyrin protein fold that have been obtained from high complexity libraries^21^ have similar size (14-18kDa) and also present a concave binding surface. 5HCS binders are concave over a larger surface area spanning the length of the protein (similar to a cupped hand), while DARPINS have long structured loops that form the majority of the binding interfaces, and are concave in a small area between these loops and adjacent helix (similar to a hand with crimped fingers); because of these structural differences the binding modes of the two with their targets are quite different (Fig. S14).

## Discussion

Our method for computationally designing small but concave proteins to bind to convex protein target sites expands the space of the protein surfaces that can be targeted by *de novo* protein design. Despite their relatively small size (120 residues), the 5HCS scaffolds span a wide range of concave shapes and have high stability (Fig. 2b, Table 1). The designed surfaces are more extensive and more concave than those obtained using our previous mini-protein binder approach: the curvature reaches –0.067 while minibinders range from –0.012 to 0.073 (more negative indicates greater curvature) and the largest distance between target bound hot-spot residues is 33 Å while minibinders range from 15 Å to 20 Å. Because of this, the 5HCS H1, H3, and H5 interface helices interact with hydrophobic pockets and patches on the target surface in ways not possible with 50-65 residue miniprotein scaffolds in which the secondary structure elements at the interface are necessarily all very close together(Fig. S13). As illustrated by the binders designed to TGFβRII and PD-L1, the dense and extended networks of hydrogen bonding residues that the 5HCS designs can present are able to satisfy the hydrogen bonding requirements of exposed target beta-strand backbone polar atoms, which enables binding modes which span both sides of the beta sheet; this is almost impossible to achieve with smaller miniproteins. The 5HCS binders can interact with beta-stands either parallelly using helix H3 with H1 and H5 flanking the sheet (5HCS_TGBR2_1) or perpendicularly with sides chains from all H1, H3 and H5 (5HCS_PDL1_1).

**Table 1.** Physicochemical properties and interface profiles of the optimized de novo 5HCS binders.

| Target | Binder ID | K<sub>D</sub> (nM) | T<sub>M</sub> (°C) | Buried surface area<br>polar / apolar (Å<sup>2</sup>) | Convexity<br>binder / target (1/Å) |
| --- | --- | --- | --- | --- | --- |
| TGFβRII | 5HCS_TGFBR2_1 | < 1 | > 95 | 637.6 / 1043.2 | -0.0669 / 0.056 |
| CTLA-4 | 5HCS_CTLA4_1 | < 0.1 | > 95 | 595.6 / 1266.1 | -0.0593 / 0.058 |
| PD-L1 | 5HCS_PDL1_1 | 0.646 ± 0.02 | > 95 | 710.4 / 1108.9 | -0.0310 / 0.001 |

The high affinity and potent signaling pathway modulation possible with the TGFβRII, PD-L1, and CTLA-4 convex binders described here demonstrates the considerable potential of our approach for targeting critical cell surface receptors. The current designs provide new routes for manipulating signaling and checkpoint blockade to be explored in future in vivo studies, and more generally our approach considerably expands the scope of de novo binder design.

## Methods and Protocols

### 5HCS Scaffolds Library Design

#### Backbone generation

The backbones were designed by taking a library of loops and helices drawn from previous successful mini-proteins and assembling them into helix-turn-helix-turn modules of 30-50 amino acids. The modules were then repeated 3 times to give a repeat protein. All possibilities of N– and C– terminal truncation were assessed and the most concave compact structure under 120 amino acids was chosen. The backbones were diversified using the Rosetta HybrizeMover using the backbones themselves as templates.

#### Sequence design and filtering

The generated backbones were designed using standard Rosetta LayerDesign protocol^22^. The heavy atoms from the residues at the concave surfaces were selected by secondary structure and rosetta layerselector. RANSAC was used to fit spherical surfaces from the coordinates of the interfacial atoms with a threshold of 1 A and max iteration 100k. The algorithm was implemented by python. By definition, convexity of the surface is the reciprocal of the radius. The designed scaffolds were later filtered by the AlphaFold2 with a mean plDDT cutoff of 80 and AccNet with a mean plDDT cutoff of 0.8. There are finally 7476 scaffolds meeting all the criteria. (library availability: https://github.com/proteincraft/5HCS)

### Protein Surface Convexity Calculation

#### Protein complex structure extraction

Pairs of interacting chains were extracted from high quality crystals from PDB. The pairs of protein complex structures were filtered by interfacial profiles, including the length of each partner’s and delta solvent accessible surface area (dSASA). Then we clustered them 40% sequence identity on both chains, and selected representatives favoring higher resolution and shorter proteins.

#### Convexity Calculation

We calculated atomic SASA for both partners from protein-protein complex pairs in apo and holo structures. Heavy atoms with a difference of 0.5 A^2^ are defined as interfacial residues. RANSAC was used to fit spherical surfaces from the coordinates of the interfacial atoms with a threshold of 1 A and max iteration 100k. The algorithm was implemented by python. By definition, convexity of the surface is the reciprocal of the radius.

#### Concave and Convex Definition

To define whether the surface is concave or convex, the geometry centers of heavy atoms of the proteins and interfacial atoms were firstly calculated. The inner product of interfacial atoms centers to protein centers and interracial atoms to fitted centers was calculated. Those surfaces with minus results are defined concave, vice versa. (code availability: https://github.com/proteincraft/5HCS)

### Interface Design and Filtering

#### Docking and Interface design

For TGFβRII binder design, the 5HCS or mini protein libraries were docked to the target binding site using the previously reported method^3^. Docked poses of the 5HCS library were filtered by binding orientation. Only designs with interfacial residues as the concave surfaces were kept. Interface sequence design was performed using ProteinMPNN^15^ with target sequences fixed as native sequences as previously reported. The designs were later filtered by ddG (less than –40), contact molecular surface (larger than 400) and pAE (less than 10) from AlphaFold2 initial guess^14^. Finally, 67 and 4310 designs from 5HCS and mini protein libraries passed the filters and were tested experimentally, respectively.

For CTLA-4 binder design, the 5HCS libraries were docked to the target binding site using the previously reported method^3^. Docked poses of the 5HCS library were filtered by binding orientation. Only designs with interfacial residues as the concave surfaces were kept. Interface sequence design was performed using previously reported protocol. The designs were later filtered by ddG (less than – 40), contact molecular surface (larger than 400). Finally, 4600 designs from 5HCS passed the filters and were tested experimentally.

For PD-L1 binder design, the 5HCS libraries were docked to the target binding site using the previously reported method^3^. Docked poses of the 5HCS library were filtered by binding orientation. Only designs with interfacial residues as the concave surfaces were kept. Interface sequence design was performed using ProteinMPNN with target sequences fixed as native sequences as previously reported. The designs were later filtered by ddG (less than –40), contact molecular surface (larger than 400) and pAE (less than 10) from AlphaFold2 initial guess^14^. Finally, 96 designs from 5HCS libraries passed the filters and were tested experimentally.

#### Combinatorial Library Design

The hits screened from the initial designs were further optimized by the virtual optimization protocol. Interfacial residues were re-sampled massively (5000 replicates) using ProteinMPNN^15^ with a higher temperature of 0.4. As the binding pattern stays mostly the same, the re-sampled designs were later assessed by delta ddG predicted by AlphaFold2 initial guess^14^. Designs with lower ddG than the initial hits were aligned by primary sequences. At each residue position, the more times of one type of mutation showed up the more likely the mutation will improve the binding affinity. We then ordered Ultramer oligonucleotides (Integrated DNA Technologies) containing the degenerate codons for the mutations predicted to be beneficial. The constructed libraries were transformed into Saccharomyces cerevisiae EBY100. The transformation efficiencies were around 10^7^.

#### Yeast Surface Display

Saccharomyces cerevisiae EBY100 strain cultures were grown in C-Trp-Ura medium supplemented with 2% (w/v) glucose. For induction of expression, yeast cells were centrifuged at 4,000g for 1 min and resuspended in SGCAA medium supplemented with 0.2% (w/v) glucose at the cell density of 1 × 10^7^ cells per ml and induced at 30 °C for 16–24 h. Cells were washed with PBSF (PBS with 1% (w/v) BSA) and labeled with biotinylated targets using two labeling methods: with-avidity and without-avidity labeling. For the with-avidity method, the cells were incubated with biotinylated target, together with anti-c-Myc fluorescein isothiocyanate (FITC, Miltenyi Biotec) and streptavidin– phycoerythrin (SAPE, ThermoFisher). The concentration of SAPE in the with-avidity method was used at one-quarter of the concentration of the biotinylated targets. For the without-avidity method, the cells were first incubated with biotinylated targets, washed and secondarily labeled with SAPE and FITC. All the original libraries of de novo designs were sorted using the with-avidity method for the first few rounds of screening to exclude weak binder candidates, followed by several without-avidity sorts with different concentrations of targets. For SSM libraries, two rounds of without-avidity sorts were applied and in the third round of screening, the libraries were titrated with a series of decreasing concentrations of targets to enrich mutants with beneficial mutations. The combinatorial libraries were enriched at medium concentration of target for two rounds by collecting the top 10% of the binding population. In the third round of sorting, the enriched library was titrated to with a series of decreasing concentrations of targets. The several binding populations with lowest concentration of target were collected.

### Protein Expression and Purification

Synthetic genes were optimized for *E. coli* expression and purchased from IDT (Integrated DNA Technologies) as plasmids in pET29b vector with a TEV-cleavable hexa-histidine affinity tag. Plasmids were transformed into BL21* (DE3) *E. coli* competent cells (Invitrogen). Single colonies from agar plate with 100 mg/L kanamycin were inoculated in 50 mL of Studier autoinduction media 45, and the expression continued at 37 °C for over 24 hours. The cells were harvested by centrifugation at 4000 g for 10 min, and resuspended in a 35 mL lysis buffer of 300 mM NaCl, 25 mM Tris pH 8.0 and 1 mM PMSF. After lysis by sonication and centrifugation at 14000 g for 45 min, the supernatant was purified by Ni^2+^ immobilized metal affinity chromatography (IMAC) with Ni-NTA Superflow resins (Qiagen). Resins with bound cell lysate were washed with 10 mL (bed volume 1 mL) of washing buffer (300 mM NaCl, 25 mM Tris pH 8.0, 60 mM imidazole) and eluted with 5 mL of elution buffer (300 mM NaCl, 25 mM Tris pH 8.0, 300 mM imidazole). Both soluble fractions and full cell culture were checked by SDS-PAGE. Soluble designs were further purified by size exclusion chromatography (SEC). Concentrated samples were run in 150 mM NaCl, 25 mM Tris pH 8.0 on a Superdex 75 Increase 10/300 gel filtration column (Cytiva). SEC-purified designs were concentrated by 10K concentrators (Amicon) and quantified by UV absorbance at 280 nm.

### Biolayer Interferometry

Binding assays were performed on an OctetRED96 BLI system (ForteBio) using streptavidin-coated biosensors. Biosensors were equilibrated for at least 10 min in Octet buffer (10 mM Hepes pH 7.4, 150 mM NaCl, 3 mM EDTA, 0.05% Surfactant P20) supplemented with 1 mg/mL bovine serum albumin (SigmaAldrich). For each experiment, the biotinylated target protein was immobilized onto the biosensors by dipping the biosensors into a solution with 50 nM target protein for 200 to 500 s, followed by dipping in fresh octet buffer to establish a baseline for 200 s. Titrations were executed at 25 °C while rotating at 1,000 rpm. Association of designs to targets on the biosensor was allowed by dipping biosensors in solutions containing designed proteins diluted in octet buffer for 800 to 3600 s. After reaching equilibrium, the biosensors were dipped into fresh buffer solution in order to monitor the dissociation kinetics for 800 to 3600 s. For binding titrations, kinetic data were collected and processed using a 1:1 binding model using the data analysis software 9.1 of the manufacturer. Global kinetic fitting using three concentration data was performed for K_D_ calculations.

### Circular dichroism

Far-ultraviolet circular dichroism measurements were carried out with a JASCO-1500 instrument equipped with a temperature-controlled multi-cell holder. Wavelength scans were measured from 260 to 190 nm at 25 and 95 °C and again at 25 °C after fast refolding (about 5 min). Temperature melts monitored the dichroism signal at 222 nm in steps of 2 °C min–1 with 30 s of equilibration time. Wavelength scans and temperature melts were performed using 0.3 mg ml^−1^ protein in PBS buffer (20 mM NaPO_4_, 150 mM NaCl, pH 7.4) with a 1 mm path-length cuvette.

### Cell assays

#### TGF-β luciferase reporter assay

The TGF-β inhibition assays utilizing HEK-293 cells stably transfected with the CAGA_12_ TGF-β reporter^23^ were performed as previously described^24^. Cells were maintained in DMEM containing 10% fetal bovine serum (FBS) and 1% penicillin/streptomycin. Cells were plated at 3×10^4^ cells per well in a treated 96-well plate. After 24 hours, the media was removed and replaced with fresh DMEM containing 0.1% bovine serum albumin (BSA) and a two-fold concentration series of 5HCS_TGFβR2_1. After 30 minutes, cells were stimulated with 10 pM TGF-β3. Twenty-four hours after stimulation, the cells were lysed and luciferase activity was measured using luciferin. The measurements for each condition were made in triplicate. IC_50_ values were calculated using the four parameters logistic regression by python scripts.

#### CTLA-4 blockade cell assay

The CTLA-4 Blockade Bioassay (Promega) was used as described in the product literature to compare bioacitivity of our novel high affinity CTLA-4 binders with Ipilimumab. Briefly, 25 uL of CTLA-4 effector cells prediluted into complete RPMI media supplemented with 10% FBS were added to wells of a 96-well flat-bottomed white luminescence plate (Costar). In a separate 96-well assay plate, antibodies and binding reagents to be tested were serially diluted into RPMI media at three times the intended final concentration. Activity of the CTLA-4 binders was compared to a control hIgG antibody (Biosciences) and the FDA-approved anti-CTLA-4 mAb Ipilimumab. From this 25uL of each diluted reagent was transferred to the wells containing CTLA-4 effector cells and subsequently 25uL of the aAPC/Raji Cells were also added. The resulting reactions were incubated for 16 hours at 37C in a humidified CO 2 incubator. After incubation, 75uL of prepared Bio-Glo reagent (Promega) was added to each well, incubated for 5min at room temperature with gentle shaking at 300 rpm and luminescence measured on an Envision plate reader (Perkin Elmer). The raw luminescence data was normalized using the following formula: *(RLU signal–background)/(RLU no antibody–background),* where the background and no antibody control values were each calculated from an average of three wells with no cells or cells but no antibody respectively. EC_50_ values were calculated using the four parameters logistic regression by python scripts.

#### PD-L1 blockade cell assay

The assays were performed according to manufacturer’s instructions (Promega). Briefly, PD-L1 aAPC/CHO-K1 cells were thawed in a 37 ℃ water bath until just thawed and transferred to pre-warmed media (90% Ham’s F12 / 10% FBS). Cells were mixed and immediately seeded to the inner 60 wells of a 96 well flat bottom white cell culture plates at 100 ul volume. 100 ul of media was also added to the outside wells to prevent evaporation. Cells were incubated for 16 hours in a 37 ℃, 5% CO₂ incubator. At the end of the incubation period, 95 ul of media was removed from each of the wells. Immediately after 40 ul of appropriate antibody or binder dilutions were added to individual wells. PD-1 effector cells were thawed in similar fashion as for PD-L1 aAPC/CHO-K1 cells and transferred to pre-warmed assay buffer (99% RPMI 1640 / 1% FBS). 40 ul of PD-1 effector cells were added to the inner 60 wells of the assay plate. 80 ul of assay buffer was added to outside wells to prevent evaporation. The assay plate was incubated for 6 hours in a 37 ℃, 5% CO₂ incubator. At the end of incubation plates were removed from the incubator and equilibrated to ambient temperature (22∼25 ℃). 80 ul of Bio-Glo reagent was added to each well and incubated for 10 mins. Luminescence was measured using the BioTek Synergy Neo2 multi-mode reader. EC_50_ values were calculated using the four parameters logistic regression by python scripts.

### Specificity Determination

#### Cell surface receptor knockouts

A431, Jurkat, and HEK293T cells had PD-L1, CTLA-4, or TGF-B knocked out respectively via CRISPR RNP transfection. RNP complexes were formed by incubating 4 ul of 80 uM guide RNA (IDT guides: Hs.Cas 9.CD274.1.AA, Hs.Cas 9.CD274.1.AB, Hs.Cas 9.CTLA4.1.AA, Hs.Cas 9.CTLA4.1.AB, Hs.Cas9.TGFBR2.1.AA, Hs.Cas 9.TGFBR2.1.AB) with 4 ul of 80 uM tracrRNA (IDT cat. 1072533) at 37°C for 30 minutes. To generate complete RNPs, 4 ul of 40 uM guide complex was incubated with 4 ul of 40 uM cas9-NLS (Berkeley MacroLab) at 37°C for 30 minutes. For electroporation, 2×105 cells of each cell type in 20 ul of electroporation buffer (Lonza, cell line SF for A431 and HEK293T, cell line SE for Jurkat) were mixed with 1 ul of electroporation enhancer (IDT cat. 1075916) and 2 ul of assembled RNPs prior to loading 20 ul into an electroporation cuvette strip (Lonza cat. V4XC-2032). Cells were electroporated with appropriate settings (A431:EQ-100, Jurkat:CL-120, HEK293T:DG-130). Cells were immediately rescued with warm complete media and transferred to a 24 well plate to grow after resting for 5 minutes at 37°C with 5% CO2. Cells were tested for knockout efficiency by TIDE analysis. Genomic DNA was extracted with Lucigen Quickextract (Lucigen cat. QE0905T) and amplified with NEBNext high-fidelity polymerase (NEB cat. M0541S).

#### Cellular surface staining

A431, Jurkat, and HEK293T cells were respectively stained with PD-L1, CTLA-4, or TGFβRII binder or antibody to compare specificity of de novo binders to commercial antibodies. For staining, 5×10^5^ cells were washed twice with 200 ul cell staining buffer (Biolegend cat. 420201) in a 96 well u-bottom plate. Cells were then resuspended in 50 ul of staining mixture (cell staining buffer and fluorophore-conjugated binder or antibody (Biolegend cat. 329713, 369605, 399709) and incubated on ice in the dark for 30 minutes.

Cells were washed three times with 200 ul staining buffer and then analyzed on a ThermoFisher Attune.

### Structure Determination

#### Expression and Purification

The coding sequence for residues 46-155 of human TβRII (UniProt P37173) was inserted into plasmid pET32a (EMD-Millipore) between the NdeI and HindIII sites without inclusion of any expression tags, transformed into chemically-competent *E. coli* BL21(DE3) (EMD-Millipore), expressed at 37 °C in the form of insoluble inclusion bodies, and refolded and purified to homogeneity as previously described^24^. The 5HCS_TGFBR2_1 used for crystallization was prepared as described above, followed by digestion for 12 h at 25°C with TEV protease (1:25 mass ratio) in 25 mM Tris, 100 mM Tris, pH 8.0, 1 mM DTT, 1 mM EDTA. Identity of the isolated protein products was verified by measuring their intact masses, which were found to be within 0.5 Da of the calculated masses (Thermo UltiMate UHPLC coupled to Bruker Compact QqTOF ESI quadrupole TOF mass spectrometer). The TbRII:5HCS_TGFBR2_1 complex was isolated by size exclusion chromatography using a HiLoad Superdex 75 26/60 column (GE Healthcare, Piscataway, NJ) in 25mM HEPES pH 7.5, 100 mM NaCl at a 1:1.1 ratio, with 5HCS_TGFBR2_1 being in slight excess. The complex peak fractions were pooled and concentrated to 33 mg/mL for crystal screening.

For large-scale purifications of the CTLA-4 and PD-L1 binders for crystallization, 2-liter bacterial cultures were grown in Super Broth (Teknova) media supplemented with antibiotics and antifoam 204 (Sigma) at 37 °C in LEX 48 airlift bioreactors (Epiphyte3, Canada) to an A600 of 3. The temperature was then reduced to 22 °C, isopropyl-β-D-thiogalactoside (IPTG) was added to 0.5 mM, and the cultures were incubated overnight. Cells were harvested by centrifugation at 14,000 x g and suspended in buffer containing 20 mM HEPES (pH 7.5), 500 mM NaCl, 20 mM imidazole, 0.1% IGEPAL, 20% sucrose, 1 mM β-mercaptoethanol (BME). Cells were disrupted by sonication and debris was removed by centrifugation at 45,000 x g. The supernatants were applied to a chromatography column packed with 10 ml His60 SuperFlow resin (Clontech Laboratories) that had been equilibrated with buffer A (50 mM HEPES pH 7.5, 30 mM imidazole, 500 mM NaCl, and 1 mM BME). The columns were washed with buffer A and the His_6_-binder proteins were eluted with buffer B (20 mM HEPES, pH 7.5, 350 mM NaCl, 400 mM imidazole, and 1 mM BME). The His_6_ tags were removed by overnight digestion at 4 °C with the TEV protease at a 1500:1 ratio of binder:TEV. The tag-free binders were then separated from His_6_-tags by Superdex 200 gel filtration equilibrated with a buffer containing 20 mM HEPES pH 7.5, 350 mM NaCl. The CTLA-4 and PD-L1 binders migrated through gel filtration as discrete peaks with estimated molecular weights of 14 kDa and 12 kDa, respectively, indicating that they are monomers in solution. The purity of the binders was judged by SDS-PAGE and Coomassie blue staining. The peak fractions from the gel filtration step were pooled and concentrated to 20-25 mg/ml in a buffer containing 20 mM HEPES (pH 7.5) and 150 mM NaCl. 5HCS_CTLA4_1cb:CTLA-4 complex were purified using size exclusion chromatography (Superdex S200) equilibrated with a buffer containing 20 mM HEPES pH 7.5, 150 mM NaCl. The peak fractions were pooled and concentrated to 7.5 mg/ml The preparations were flash frozen in liquid nitrogen and stored at –80 °C for long-term storage.

#### Protein crystallization and crystal harvesting

Crystals of the TβRII:5HCS_TGFBR2_1 complex were formed using hanging drop vapor diffusion in 24-well plates with 300 μL of well solution and siliconized glass cover slips. Crystals formed in 1-2 days at 25 °C with drops prepared by mixing 0.4 μL 25 mg/mL protein complex and 0.4 μL of 20% (w/v) PEG-MME 5K, 0.4 M (NH_4_)_2_ SO_4_, 0.1 M Tris pH 7.4, and 16 – 32 % glycerol. The crystals were mounted in nylon loops without additional cryoprotectants and with excess well solution wicked off.

Screening of 5HCS_CTLA4_1cb2 and 5HCS_PDL1_1 for crystal formation was performed using 0.8 uL (protein: reservoir solution=1:1) crystallization drops at a concentration of 15 mg/ml with a Crystal Gryphon (Art Robbins Instruments) robot, using MCSG (Microlytic), Index HT, Crystal Screen HT, and Peg Ion HT sparse matrix crystallization suites (Hampton Research). Initial crystals obtained from the sparse matrix screening were further optimized with several rounds of grid screening using a Formulator robot (Art Robbins Instruments). Optimized crystallization conditions for diffraction quality binder protein crystals and cryo-protectants used during crystal harvesting are summarized in Table S2.

#### Data collection and processing, structure refinement and analysis

The diffraction data for the TβRII:5HCS_TGFBR2_1 complex was collected at the Southeast Regional Collaborative Access Team (SER-CAT) 22-ID beamline at the Advanced Photon Source, Argonne National Laboratory. The data was integrated with XDS^25^ and the space group (P2_1_2_1_2_1_ with dimensions a,b,c = 47.98 Å, 57.17 Å, 78.80 Å and α,β,γ = 90°, 90°, 90°) was confirmed via pointless^26^. The data was reduced with aimless^27^, ctruncate^28–32^ and the uniquify script in the CCP4 software suite^33^. Phasing was performed with Phaser^34^, initially with the 1.1 Å TβRII X-ray structure (PDB 1M9Z), followed by the predicted 5HCS_TGFBR2_1 structure. Several cycles of refinement using Refmac5^35–42^ and model building using COOT^43^ were performed to determine the final structure. Data collection and refinement statistics are shown in Table S2.

Data from the crystals of CTLA-4 binder were collected on a Dectris Pilatus 6M detector, with a wavelength of 0.98 Å, on the ID-31 (LRL-CAT) beamline at the Argonne National Laboratory (Table S2). Single crystal data were integrated and scaled using iMosflm^44^ and aimless^33^, respectively. Diffraction was consistent with the orthorhombic space group P2_1_2_1_2_1_ (unit cell dimensions are in Table S2) and extended to 1.85 Å resolution with one molecule (chain A) in the asymmetric unit. Data for the PD-L1 binder crystals were collected on a Dectris EIGER X 9M detector, with a wavelength of 0.92 Å, on the 17-ID-1 (AMX) beamline at the Brookhaven National Laboratory (Table S2). Data for the CTLA-4-binder complex crystals were collected on a Dectris EIGER X 9M detector, with a wavelength of 0.98 Å, on the 17-ID-2 (FMX) beamline at the Brookhaven National Laboratory (Table S2). The datasets were indexed, integrated, and scaled using fastDP, XDS^25^ and aimless^44^, respectively. The PD-L1 crystals belong to tetragonal space group and diffracted to 1.88 Å. The CTLA-4-binder complex crystal belongs to C2 space group and diffracted to 2.72 Å. Initial phases of 5HCS_CTLA4_1cb2, 5HCS_PDL1_1 and 5HCS_CTLA4_1cb:CTLA-4 complex were determined by molecular replacement (MR) with Phaser^34^. using coordinates of the computationally designed respective binders and binder complex; the initial MR coordinate was manually inspected and corrected using Coot^43^. The model was refined with Phenix-Refine^45^. Analyses of the structures were performed in Coot and evaluated using MolProbity^46^; B-factors were calculated using Baverage program in CCP4 suite^47^. The crystallographic model exhibited excellent geometry with no residues in disallowed regions of the Ramachandran plot^48^.

Crystallographic statistics and RCSB accession codes are provided in Table S2. All figures depicting structure were generated with PyMol, unless stated otherwise.

## Acknowledgement

Funding for this work was provided by a gift from Gates Ventures (W.Y., L.S., D.B.), the National Institute on Aging (NIA) Grants R01AG063845 (I.G., D.B.) and U19AG065156 (D.R.H.), the Audacious Project at the Institute for Protein Design (A.A., S.F.H., D.L., C.J.K., S.R.G., D.B.), grants from DARPA supporting the Harnessing Enzymatic Activity for Lifesaving Remedies (HEALR) program (HR001120S0052 contract HR0011-21-2-0012, A.K.B., B.H., D.B.) and the Synergistic Discovery and Design project (HR001117S0003 contract FA8750-17-C-0219, L.C., D.B.), the Open Philanthropy Project Improving Protein Design Fund (Y.W., D.B.), and the Howard Hughes Medical Institute (B.C., D.B.).

## Author Contributions

W.Y., D.B. designed the research. W.Y., B.C., D.H. designed the 5HCS scaffold library. W.Y., D.H., B. H. and L.C. designed the binders. W.Y., D.H., I.G., S.H. and A.A. performed library preparation, the yeast screening, expression and binding experiments. A.G. solved structure of bound and unbound CTLA-4 binder. A.G., A.H. solved the structure of the unbound PD-L1 binder. T.A.S and A.H solved the structure of bound TGFbRII binder. W.Y., D.H., Z.L., S.G., P. H. and A. M. expressed and purified proteins. W.Y. and Y.W. performed circular dichroism measurements. C.H., D.L. performed TGFb3 inhibition assay. S.G., C.K. performed CTLA-4 activation assay. C.K. performed PD-L1 activation assay. W.Y., D.H. and S.R. performed binding assays. T. S. and N. E. designs oligomer scaffolds. All authors analyzed data. L.S., D.B. supervised research. W.Y., D.B. wrote the manuscript. All authors revised the manuscript.

