## supplementary for "Design of High Affinity Binders to Convex Protein Target Sites"

### Supplementary Information

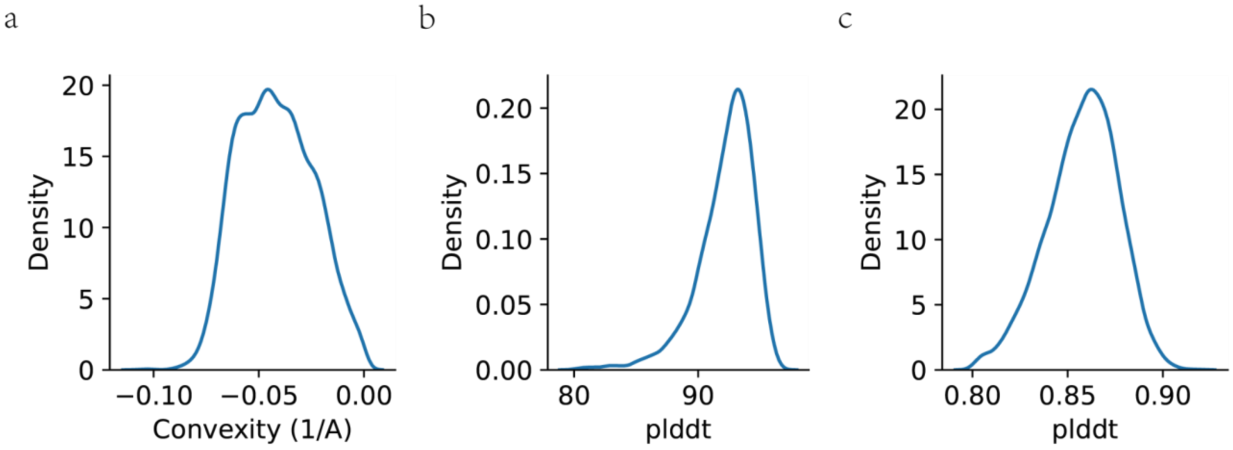


**Figure S1. Computational Characterization of 5HCS scaffolds library. a,** Distribution of curvatures of the 5HCS library. **b,** Distribution of pLDDT predicted by Alphafold2[1]. **c,** Distribution of plDDT predicted by DeepAccNet[2].

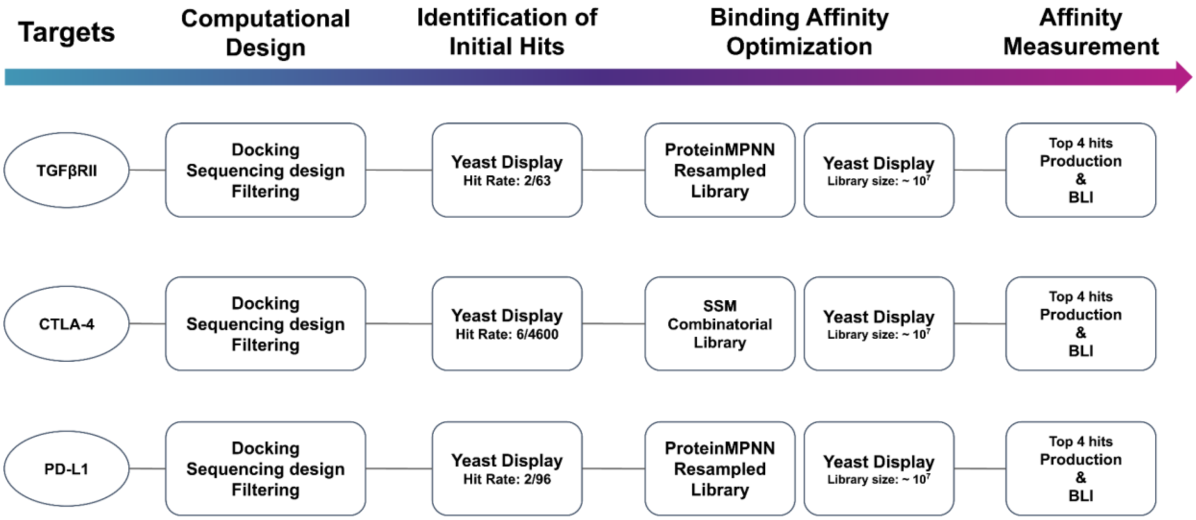


**Figure S3. Workflow of Design and Optimization of the 5HCS binders.** There are generally four stages for each target including, computational design, identification of initial hits by yeast display (first round of yeast display), binding affinity optimization ( second round of yeast display) and final affinity measurement. Library size and hit rates are labeled in the corresponding boxes.

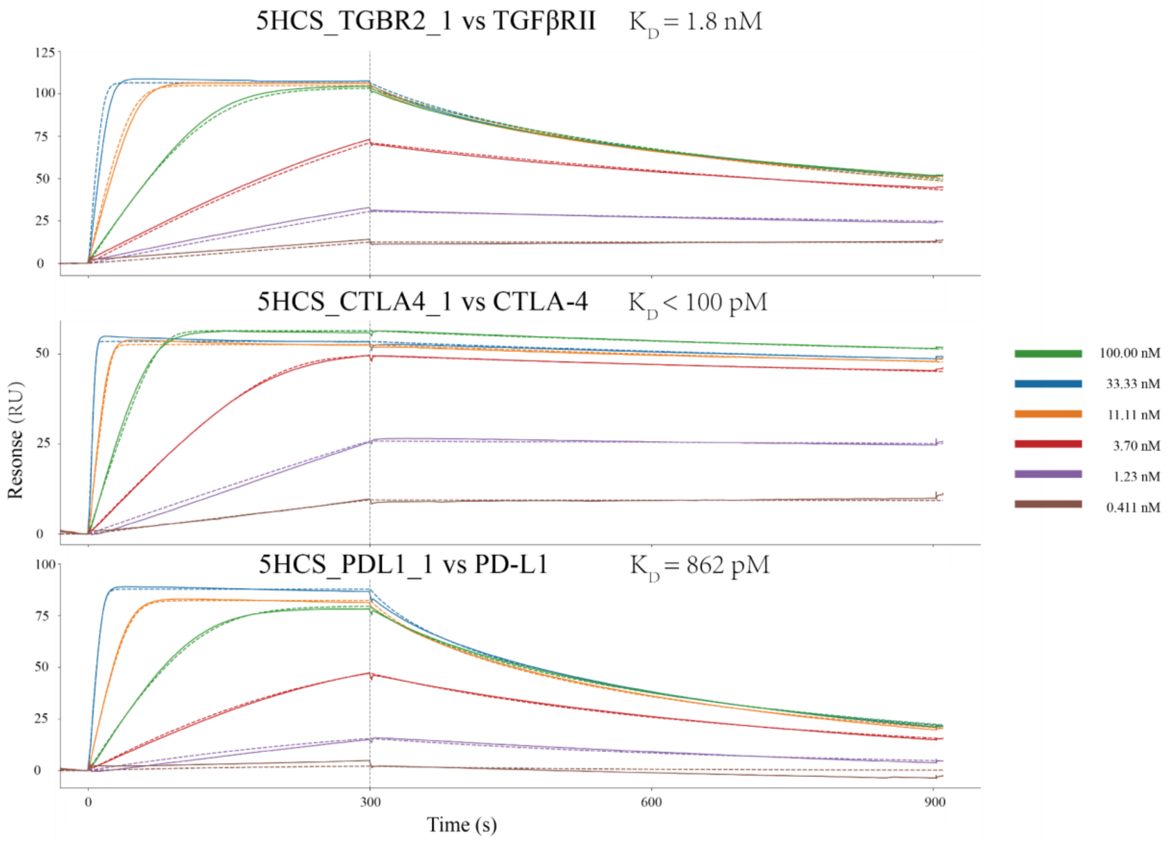


**Figure S5. Binding of 5HCS binders to corresponding targets.** Binding of 5HCS binder to corresponding targets were measured by Biacore 8K using HEPES running buffer with P20 (0.01 M HEPES pH 7.4, 0.15 M NaCl, 3 mM EDTA, 0.005% v/v Surfactant P20). Target proteins were immobilized on chips(biotin capture chip:TGFβRII, CTLA-4; protein A chip: PD-L1) Binding traces (solid lines) are fitted (dased lines) using Langumuir 1:1 interaction model for kinetic evaluation.

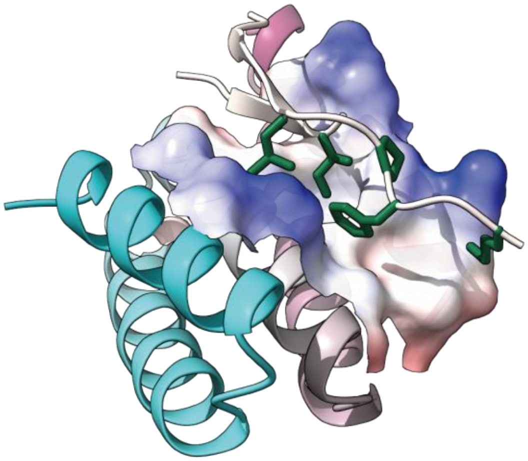

**Figure S7. De novo designed binding groove on 5HCS_TGFBR2_1 .** 5HCS_TGFBR2_1 shown in cartoon (rainbow) and electrostatic potential surface. TGFβRII binding motif shown as white cartoon and green sticks.

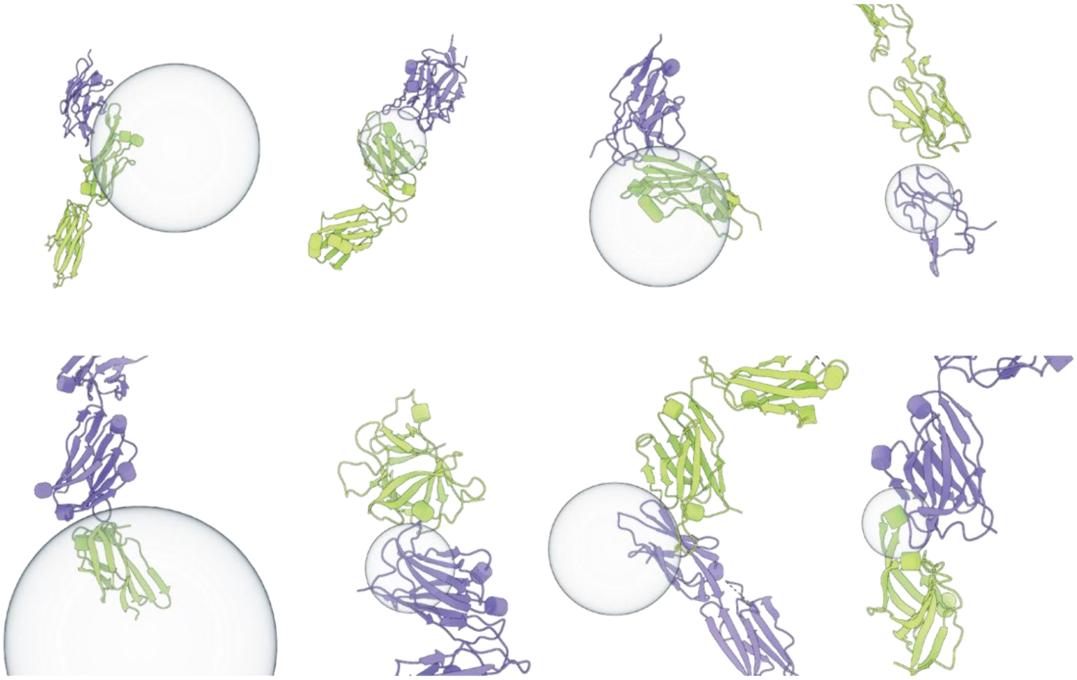

**Figure S8. Examples of Immunoglobulin domain head-to-head interactions.** PDB ID: 1i8l, 1iqd, 2jjs, 3r08, 3s35, 4g6m, 4hcr, 5d1q. Convex protein surfaces were fitted as spheres.

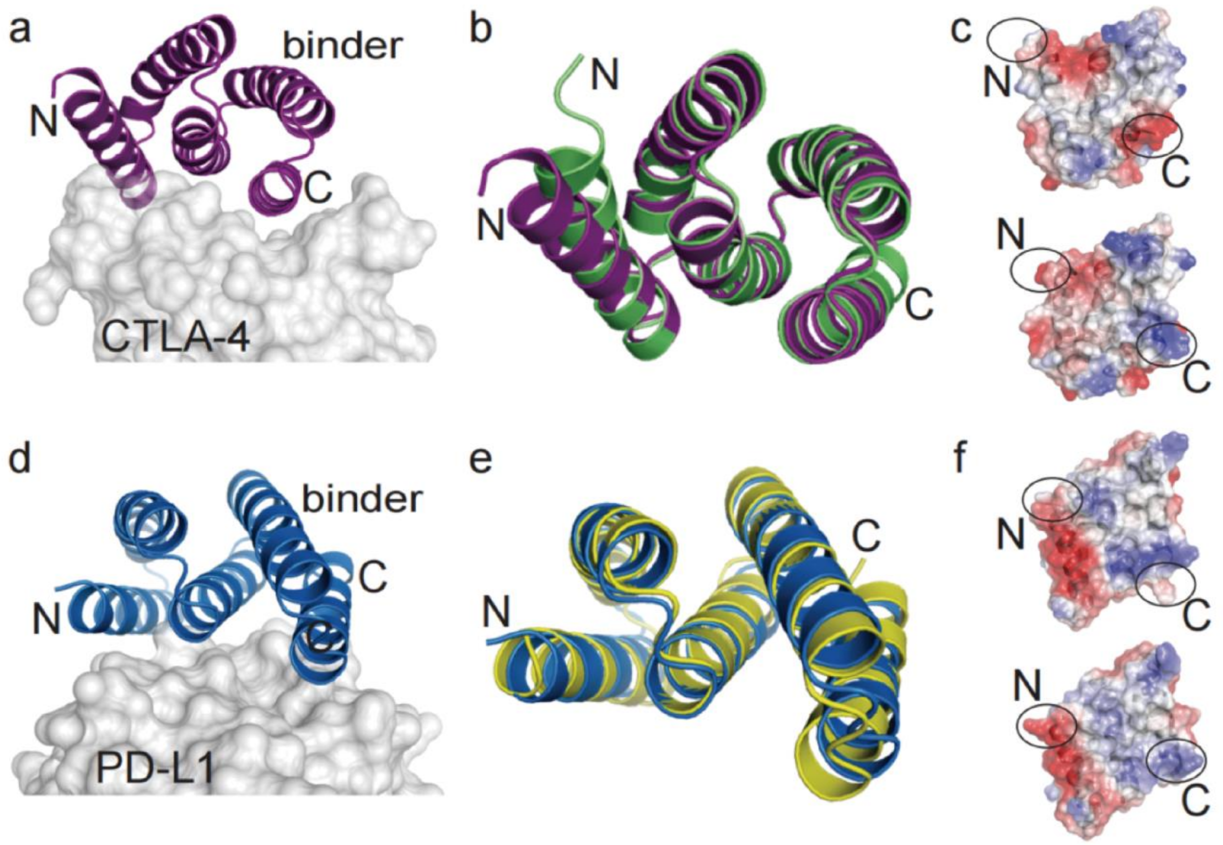


**Figure S10. High-resolution structures of 5HCS_CTLA4_2 and 5HCS_PDL1_1.**

**a**, A view of the structure of 5HCS_CTLA4_2 shown in ribbon representation (colored purple). N and C denote the location of the N and C termini. Transparent molecular surface (light gray) depicts modeled CTLA-4 to highlight the binding interface. **b**. Overlay of 5HCS_CTLA4_2 structure to the design model (green). **c**. Comparison of charge distribution at the binder interface between the crystal structure (top) and the model (bottom). **d**. A view of the structure of 5HCS_PDL1_1 (blue, ribbon representation). Part of the modeled PD-L1 is shown as a transparent molecular surface (light gray) **e.** Superimposition of 5HCS_PDL1_1 structure to the design model (yellow). **f.** Electrostatic potential at 5HCS_PDL1_1 interface (top, crystal structure; bottom, de novo model).

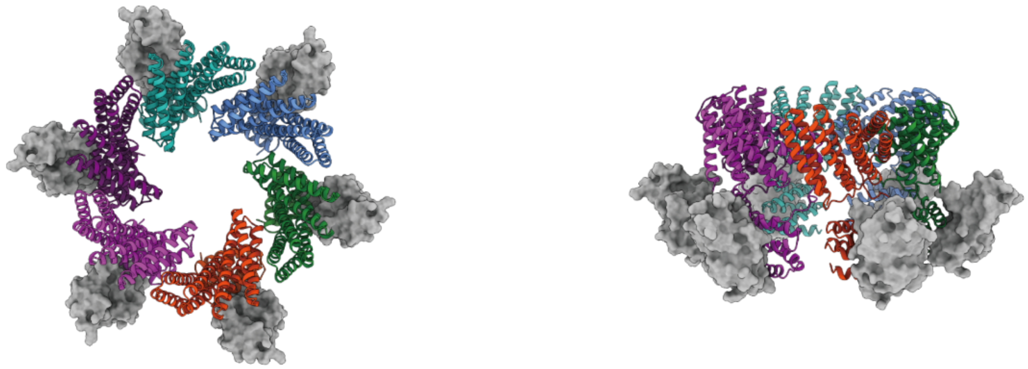

**Figure S11. 5HCS_CTLA4_1 fused with hexamers.** Top view (left) and side view (right) of the 5HCS_CTLA4_1 fusion with hexamers (cartoon) in complex with CTLA-4 (grey surface). Each chain is colored by a different color.

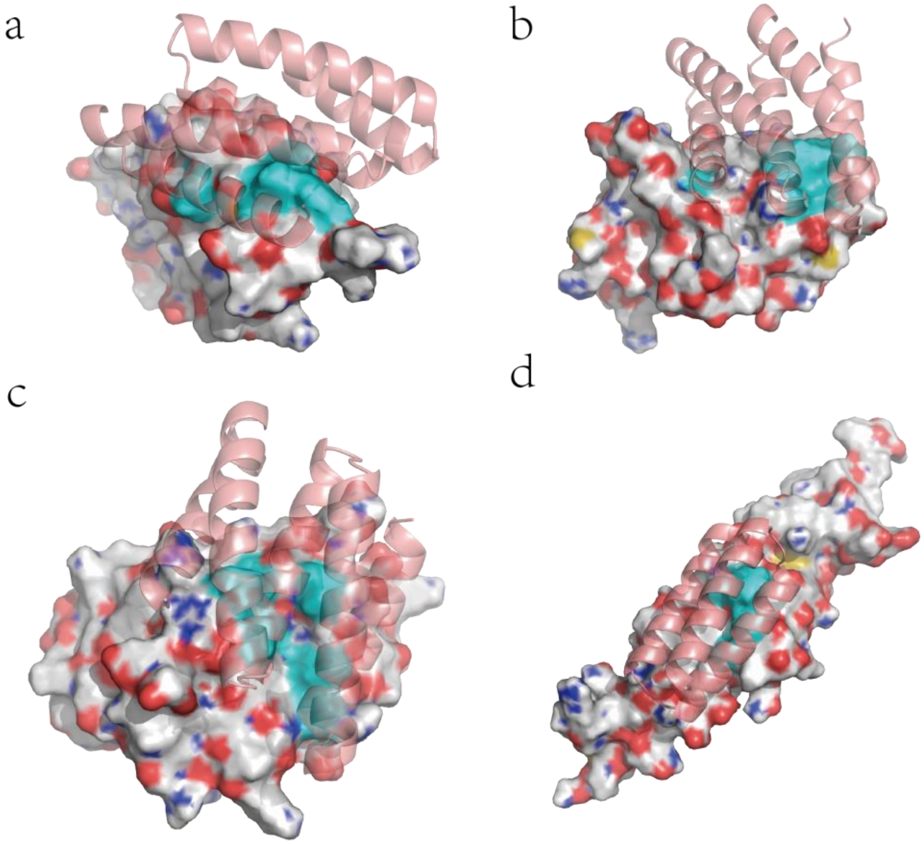


**Figure S13. Comparison of binding patches of 5HCS and minibinders.**

5HCS_TGFBR2_1 (a), 5HCS_CTLA4_1(b), 5HCS_PDL1_1(b), and Her2 minibinder. Target proteins are shown as surface and binders are shown as cartoons. Key interactions patches are colored in cyan.

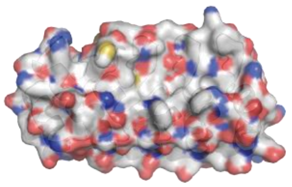

**Figure S14. Example of binding surface of Darpin (PDB ID: 8P9E).**

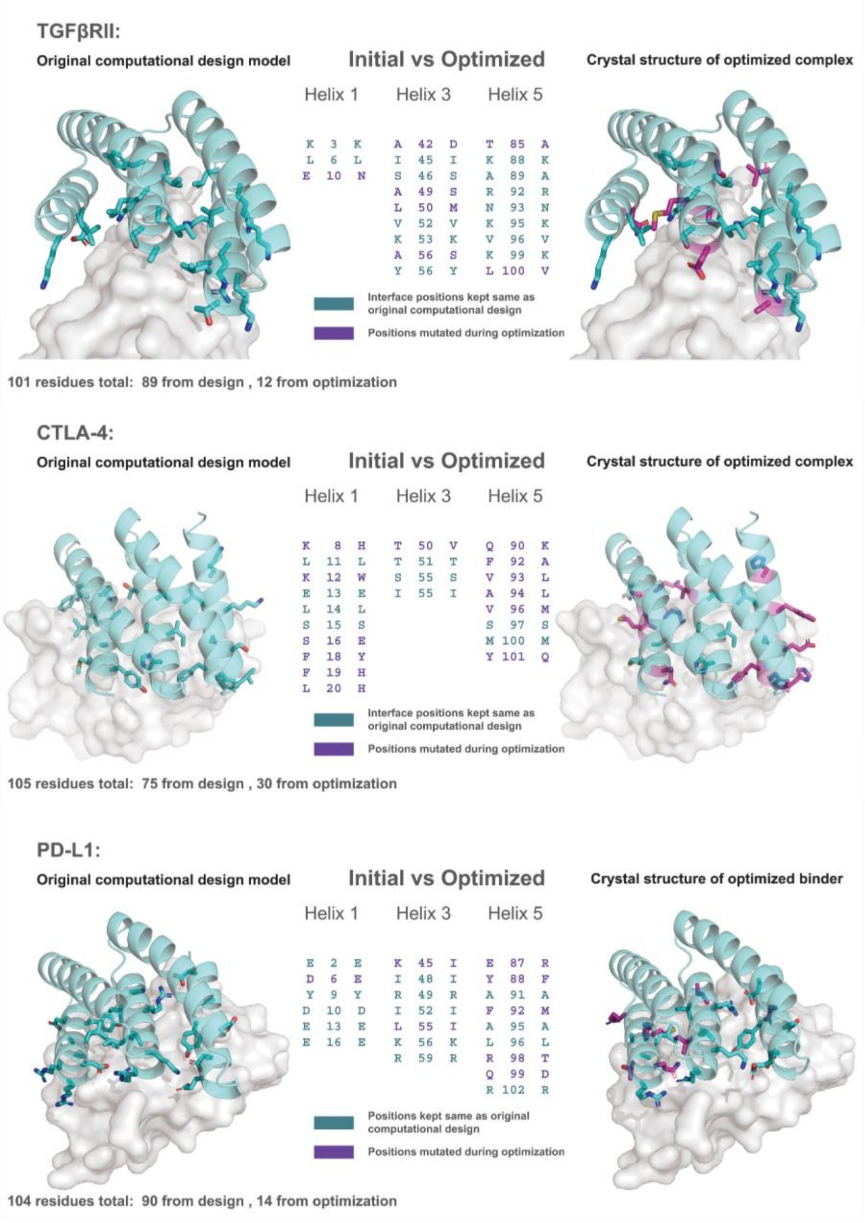


**Figure S15. Comparison of original computational design models and crystal structures.** The backbones and docking poses of originally computationally designed 5HCS binders for three targets (left) are nearly identical to the crystals of optimized 5HCS binders. A small fraction of residues were optimized through experimental selection methods. Mutations introduced on the interfaces are highlighted in the middle.

**Table S1: Interface profiles of de novo designed miniprotein binders**

| Target | Buried surface area polar/ apolar (Å$^2$) | Convexity binder / target (1/Å) |
|---|---|---|
| EGFR | 587.5 / 1150.5 | 3.25e-2 / -2.08e-2 |
| FGFR2 | 565.34 / 1394.2 | -1.5e-3 / -4.6e-2 |
| H3 | 740.3 / 892.5 | 1.48e-2 / -3.08e-4 |
| IGF1R | 1064.9 / 542.2 | 7.31e-2 / -8.96e-3 |
| IL7Ra | 763.8 / 1039.6 | 1.70e-2 / 9.46e-3 |
| InsulinR | 482.9 / 1101.8 | -1.16e-2 / 1.99e-2 |
| PDGFR | 687.7 / 1398.0 | 2.75e-2 / -2.89e-2 |
| TGFb | 861.0 / 1026.3 | -4.87e-4 / -6.08e-3 |
| Tie2 | 583.8 / 1032.7 | 8.05e-4 / 1.06e-2 |
| TrkA | 438.2 / 1161.2 | 5.17e-2 / 1.29e-2 |
| VirB8 | 413.1 / 1293.4 | 4.99e-2 / -8.9e-3 |

**Table S2: Crystallization conditions and Crystallographic data statistics**

| **Protein** | **TGFβRII binder (5HCS_TGBR2_1) complex** | **CTLA-4 binder (5HCS_CTLA4_1) complex** | **CTLA-4 binder (5HCS_CTLA4_2)** | **PD-L1 binder (5HCS_PDL1_1)** |
|---|---|---|---|---|
| **Ligand[$]** | **none** | **none** | **none** | **Indole** |
| **Crystallization condition** | 20% (w/v) PEG-MME 5K, 0.4 M $(NH_4)_2$ $SO_4$, 0.1 M Tris pH 7.4, and 16 – 32 % glycerol | 22% (w/v) PEG 3350 and 0.2 M KCl | 30 % (w/v) PEG 3350, 0.22 M $MgCl_2$ and 0.0.8 M Bis-Tris pH 5.5 and 0.15 M HEPES pH 7.5 | 22.5% (w/v) PEG 3350, 0.08 M Bis-Tris pH 5.5 and 2% (v/v) Ethylene glycol |
| **Cryo preservation** | 16 - 32% glycerol | 8% (v/v) Ethylene glycol | 5% (v/v) Ethylene glycol | 8% (v/v) Ethylene glycol |
| **Data Collection*** | | | | |
| Source | APS 22-ID | BNL 17-ID-2 | APS 31-ID | BNL 17-ID-1 |
| Wavelength (Å) | 1.00 | 0.98 | 0.98 | 0.92 |
| Number of crystals | 1 | 1 | 1 | 1 |
| Space group | $P2_12_12_1$ | C2 | $P2_12_12_1$ | $P4_32_12$ |
| Cell dimensions | | | | |
| a,b,c (Å) | 47.98, 57.17, 78.80 | 175.75, 33.6,74.47; β = 101.15 | 39.9, 52.1, 53.5 | 67.24, 67.24, 101.9 |
| Resolution (Å) | 46.28-1.24(1.27-1.24) | 20-2.72 (2.82-2.72) | 20-1.85 (1.92-1.85) | 20-1.88 (1.95-1.88) |
| Completeness (%) | 95.7(53.4) | 99.8 (99.3) | 99.6 (99.9) | 99.8 (99.3) |
| Total reflections | 547697 | 44220 | 105488 | 286708 |

| | | | | |
|---|---|---|---|---|
| Unique reflections | 61539 | 11876 | 9930 | 19508 |
| Wilson B-factor | 14.1 | 55.07 | 26.4 | 36.85 |
| Multiplicity | 8.9(2.3) | 3.7 (3.6) | 10.6 (10.9) | 14.7 (14.9) |
| $R_{merge}$ (%) | 5.2(89.8) | 8.7 (89.7) | 8 (80.3) | 5.9 (84.7) |
| $CC_{1/2}$ (%) | 100(42.5) | 99.9 (87.2) | 99.9 (84.4) | 99.9 (89.1) |
| CC* (%) | 100(77.2) | 99.6 (87.6) | 100 (96.9) | 100 (97.1) |
| <I>/s(I) | 18.9(0.6) | 10.35 (2.08) | 18.4 (3.4) | 30.24 (3.66) |
| **Refinement*** | | | | |
| Resolution (Å) | 46.28-1.24 | 19.5-2.72 ( 2.82 - 2.72) | 19.9-1.85 (2.12-1.85) | 19.62-1.88 (1.95-1.88) |
| Reflections: work/free | 61539/3078 | 11858 (1170)/548 (67) | 9928 (985)/430 (45) | 19506 (1888)/983 (90) |
| $R_{work}$/$R_{free}$ (%) | 18.13/19.58 | 28.7 (32.6)/32.9 (37.4) | 19.1 (23.4)/21.8 (26.7) | 18.9 (25.7)/21.2 (29.9) |
| Number of TLS groups | 0 | 0 | 4 | 8 |
| Number of atoms | | | | |
| Protein | 3517 | 3482 | 2200 | 1632 |
| Ligand | 5 | 0 | 4 | 9 |
| Water | 201 | 0 | 63 | 103 |
| Average B-factors (Å$^2$) | | | | |

| | | | | |
|---|---|---|---|---|
| Protein | 20.92 | 50.3 | 29.1 | 43.23 |
| Ligand | 30.0 | | 39.6 | 51.75 |
| Water | 34.96 | | 38.9 | 49.93 |
| rmsd. | | | | |
| Bond lengths (Å) | 0.0198 | 0.004 | 0.01 | 0.007 |
| Bond angles (°) | 1.9599 | 0.73 | 1.1 | 0.88 |
| MolProbity† | | | | |
| Favored | 98.6% (217 aa) | 98% (394 aa) | 100% (98 aa) | 99.5% (208 aa) |
| Allowed | 100% (220 aa) | 100% (402 aa) | none | 100% (209 aa) |
| Outliers | none | none | none | none |
| Clash score | 100th percentile | 98th percentile | 100th percentile | 100th percentile |
| Molprobity score | 100th percentile | 98th percentile | 100th percentile | 100th percentile |
| **RCSB ID** | 8G4K | 8GAB | 8GAC | 8GAD |

* Statistics calculated using Phenix (Adam et al., 2010); highest resolution shells indicated in parentheses

† Calculated with the program MolProbity (Chen et al., 2010)
